# Expression of c-Fos and Arc in hippocampal region CA1 marks neurons that exhibit learning-related activity changes

**DOI:** 10.1101/644526

**Authors:** David Mahringer, Anders V. Petersen, Aris Fiser, Hiroyuki Okuno, Haruhiko Bito, Jean-François Perrier, Georg B. Keller

## Abstract

Immediate early genes (IEGs) are transcribed in response to neural activity and necessary for many forms of plasticity. However, the dynamics of their expression during learning, as well as their relationship to neural activity, remain unclear. Here we used two-photon imaging in transgenic mice that express a GFP-tagged variant of Arc or c-Fos and a red-shifted calcium indicator to measure learning-related changes in IEG expression levels and neural activity in hippocampal region CA1 as mice learned to perform a two-alternative forced choice task. Neural activity levels correlated positively with IEG expression levels *in vivo*. In addition, we found that with learning, a subset of neurons in CA1 increased their responses to the reward-predicting cue, and IEG expression levels early in learning were selectively increased in neurons that would exhibit the strongest learning-related changes. Our findings are consistent with an interpretation of IEG expression levels as markers for experience dependent plasticity.

## INTRODUCTION

Learning is associated with persistent changes in the central nervous system. These changes can manifest as a strengthening or weakening of synaptic weights (Hebb, 1949) as they occur during long-term potentiation (LTP) and long-term depression (LTD) (Bi and Poo, 1998; Bliss et al., 1973), or the appearance or elimination of synapses (Engert and Bonhoeffer, 1999; Maletic-Savatic et al., 1999). The molecular and gene expression changes underlying this neural plasticity are not fully understood but have been shown to involve increases in the expression of a set of genes, referred to as immediate early genes (IEGs) (Okuno, 2011). Plasticity is thought to be triggered by specific changes in Ca^2+^ concentration that activate calcium-dependent kinase cascades, which in turn activate cAMP-response element binding proteins (CREB) (Mermelstein et al., 2000). This results in the upregulation of the expression of transcription factors like c-Fos (Worley et al., 1993) and other IEG products like Arc (activity-regulated cytoskeletal associated protein, or Arg 3.1; (Vazdarjanova et al., 2006)). As a consequence, the expression of *c-fos* and *Arc* are often interpreted as a marker of neural activity and plasticity (Minatohara et al., 2015). In the hippocampal formation, the induction of LTP and exposure of an animal to spatial tasks are followed by an increase in the level of mRNA of *c-fos* (Cole et al., 1989; Dragunow and Faull, 1989; Guzowski et al., 2001; Ranieri et al., 2012; Vann et al., 2000) and *Arc* (Link et al., 1995; Lyford et al., 1995). The expression of both *c-fos* and *Arc* are also involved in learning-related plasticity: A central nervous system-wide knockout of *c-fos* results in deficits in hippocampus-dependent spatial and associative learning tasks in adult mice (Fleischmann et al., 2003). *Arc* has been shown to regulate spine morphology (Peebles et al., 2010), and is critically involved in LTD of synapses (Guzowski et al., 2000; Jakkamstti et al., 2013; Plath et al., 2006; Wall et al., 2018). In addition, Arc protein has been shown to be targeted to silent synapses where it mediates AMPA receptor endocytosis and thereby induces synaptic weakening (Okuno et al. 2012). Neurons in the hippocampal formation that express *c-fos* or *Arc* have been shown to be critical for memory formation and recall: the selective activation of neurons that express *c-fos* during fear conditioning can reactivate the fear memory (Garner et al., 2012). The inhibition of these neurons selectively in CA1 suppresses the expression of the fear memory (Tanaka et al., 2014), while selective re-activation of these neurons in the dentate gyrus induces freezing-behavior (Liu et al., 2012; Ryan et al., 2015). Inhibition of neurons that express *Arc* in the dentate gyrus or in CA3 during contextual fear conditioning results in an impairment of the fear memory (Denny et al., 2014). It is still unclear, however, how IEG expression levels are dynamically regulated by neural activity during learning in the hippocampal formation. Here, we describe the expression dynamics of *c-fos* and *Arc* in CA1 pyramidal neurons during learning of a two-alternative forced choice (2AFC) tone discrimination task. We show that neurons with the highest expression of IEGs during the early phase of learning become selectively responsive to the task relevant tone cues late in learning.

## RESULTS

To measure learning-related changes in both IEG expression levels and neural activity in the same CA1 neurons we expressed a genetically encoded calcium indicator (Dana et al., 2016) by a local injection of an AAV vector (AAV2/1-EF1a-jRGECO) in mice that expressed either a c-Fos-GFP (Barth et al., 2004) or an Arc-GFP (Okuno et al., 2012) fusion protein. We then trained these mice in a 2AFC tone discrimination task. Throughout these experiments, mice were head-fixed under a two-photon microscope, held in a cylinder with two lick spouts presented in front of them, and trained to lick on one of two lick spouts (left or right) depending on the frequency of a pure tone presented to them (**Figure 1A**). Mice were accustomed to the setup and the experimenter in two sessions preceding the start of training. During these sessions, mice received water rewards in random alternation from both lick spouts to accustom them to licking on the lick spouts. Training in the 2AFC task was then spread over the subsequent 7 sessions that each lasted one hour and occurred daily. All training trials were initiated by the presentation of one of two randomly selected visual stimuli (a full-field grating) presented for 2 seconds on a toroidal screen in front of the mice (**Figure 1B**). The identity of the visual stimulus was not informative of the correct lick spout. Following the visual stimulus, one of two pure tones (6 kHz or 11 kHz) was presented for 4 s. The identity of this tone stimulus indicated the correct lick spout (left or right). All licks that occurred within 2 seconds of the tone onset were ignored and the first lick following this 2 second grace period was used to determine the choice of the mouse. Correct choices were rewarded with a drop of water. Choosing the incorrect lick spout or failure to lick in the response window (2 s to 4 s after tone onset) resulted in a mild air puff, as well as an additional delay of 5 seconds added to the inter trial interval of 14 seconds. To facilitate learning, mice received a reward on the corresponding lick spout, independent of which spout they licked on, in 10% of randomly selected trials. Mice learned to perform this task over the course of the 7 training sessions, and performance of the mice was above chance starting with day 3 of training (**Figure 1C**).

**Figure 1.**
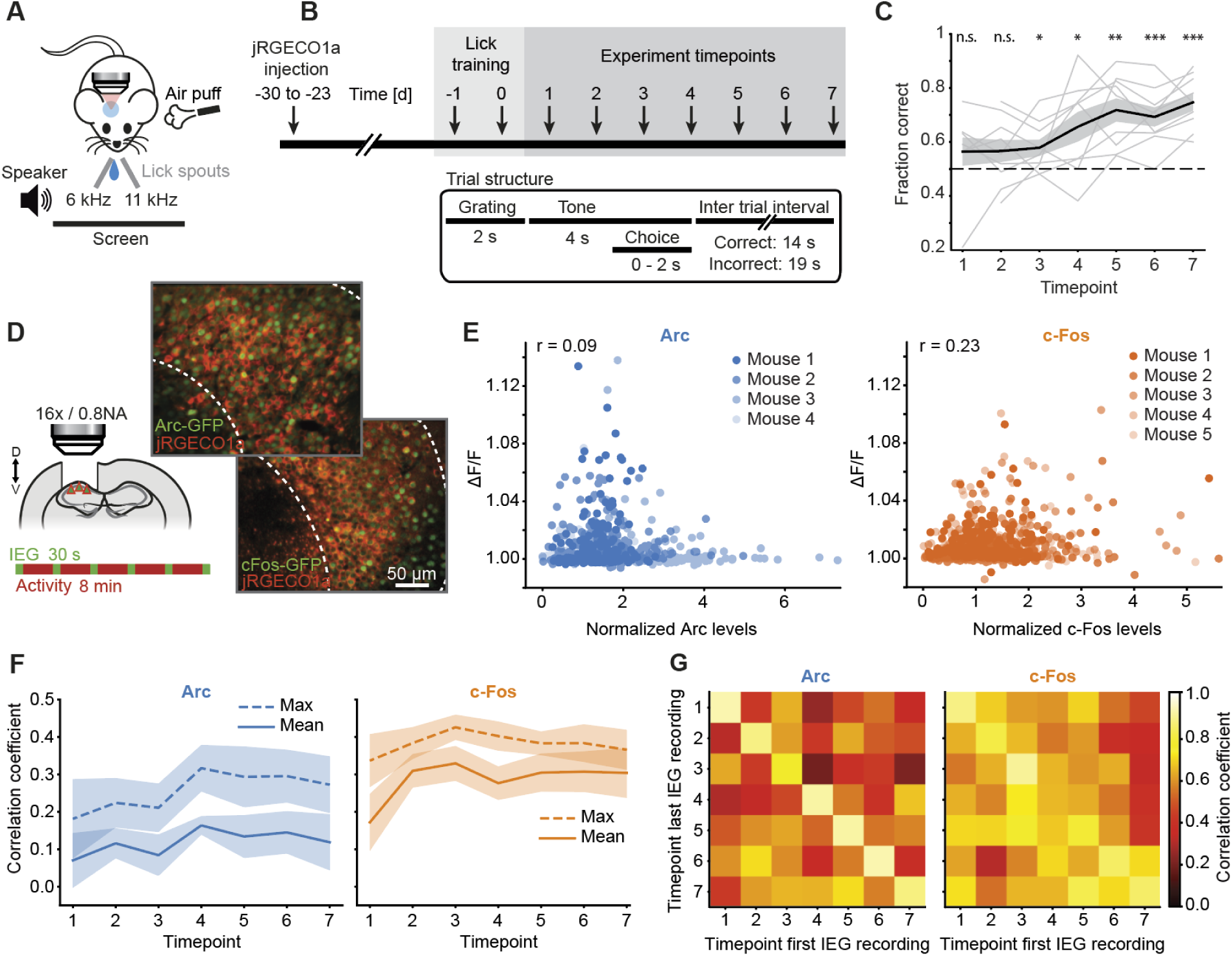
Measuring calcium activity and Arc or c-Fos expression levels chronically during learning of a two-alternative forced choice task. (**A**) Schematic of the experimental setup. Mice were head-fixed under a two-photon microscope and trained in a 2AFC tone discrimination task to lick either left or right depending on the frequency of a tone cue presented to them. Correct choices were rewarded with a drop of water, incorrect choices punished with an air puff. (**B**) Top: Schematic of the experimental timeline. Mice were exposed to the setup and familiarized with the lick spouts in two sessions preceding the experiment (lick training). Training in the task occurred in 7 sessions spaced by one day. Bottom: Schematic of the trial structure. Trial start was signaled by the presentation of one of two gratings, followed by one of two tones (6 kHz or 11 kHz) that signaled to lick right or left respectively. All licks during the 2 seconds after the tone onset were ignored. The first lick after this 2 second grace period was used to determine the choice of the mouse. Choosing the wrong lick spout or failure to lick within the response window (2 s to 4 s after tone onset) triggered an air puff and resulted in a longer inter trial interval (19 s) than following a correct choice (14 s). (**C**) Performance in the task quantified as the fraction of correct trials as a function of training day (timepoint). Data from individual mice are shown in thin gray lines, mean and SEM over all mice are indicated by the black line and gray shading (9 mice). Note, 2 mice did not perform the task on day 1. Dashed black line marks chance performance (50%). Performance is significantly above chance on timepoints 3-7 (*: p < 0.05; **: p < 0.01; ***: p < 0.001 n.s.: p > 0.05, Student’s t-test). (**D**) Left: Schematic of CA1 imaging strategy. Cortex overlying CA1 was removed and an AAV2/1-EF1a-jRGECO injected in CA1. IEG and neural activity were recorded in alternating intervals of 30 s IEG recording and 8 minutes calcium recording. Top right: Example two-photon image of CA1 neurons co-expressing the genetically encoded calcium indicator jRGECO1a (red) and the fusion protein Arc-GFP (green). Bottom right: Example two-photon image of CA1 neurons co-expressing the genetically encoded calcium indicator jRGECO1a (red) and the fusion protein c-Fos-GFP (green). Dashed white lines mark CA1 borders. (**E**) Scatter plot of average calcium activity and IEG expression levels recorded 34 to 42 minutes later for Arc (left) and c-Fos (right) averaged across all timepoints per neuron (Arc-GFP: 1271 neurons; c-Fos-GFP: 1819 neurons). Different colors mark data from different animals (4 Arc-GFP mice, 5 c-Fos-GFP mice). r-value is the average Pearson correlation coefficient across mice. (**F**) Pearson’s correlation coefficient between IEG expression with average (solid lines) or maximum (dashed lines) activity for Arc (left) and c-Fos (right) as a function of timepoint. Shading indicates SEM over mice. (**G**) Spearman’s correlation coefficient of IEG expression between the first (columns) and last (rows) IEG recording in each timepoint for Arc (left) and c-Fos (right).

Throughout all training sessions we chronically recorded IEG expression levels and neural activity in the same CA1 pyramidal neurons using two-photon imaging (1271 neurons in 4 Arc-GFP mice, 1819 neurons in 5 c-Fos-GFP mice). To access CA1 for two-photon imaging, we removed the cortex above the left or right hippocampus, injected AA2/1-EF1a-jRGECOa to express the calcium indicator in CA1 and implanted a cranial window 23 to 30 days prior to the start of training, as previously described (Fiser et al., 2016). Calcium activity was measured throughout the training sessions, while IEG expression levels were measured every 8 min during 30 second breaks in the training paradigm (**Figures 1B and 1D**). On average, c-Fos expression levels were stable, while Arc expression levels decreased over the course of learning (**Figure S1A;** Arc: p = 0.005, R^2^ = 0.26, 4 mice; c-Fos: p = 0.746, R^2^ = 0.003, 5 mice; linear trend analysis, see Methods). At the same time, average neural activity decreased between training session 1 and 2, and then remained stable for the rest of the training sessions (**Figure S1A**).

We then quantified the relationship between calcium activity and IEG expression. Consistent with previous reports (Tanaka et al., 2018; Yassin et al., 2010), we found that average calcium activity correlated positively with expression levels recorded 34 to 42 minutes later for both Arc and c-Fos (**Figure 1E**). The cross-correlation between neural activity and IEG expression levels was increased and relatively stable over a broad range of time lags between neural activity and IEG expression measurement (**Figure S1C**). Note, as we are not artificially inducing activity and quantifying IEG expression following induction, but rather looking at the cross-correlation between neural activity and IEG expression levels over extended periods of natural activity, the width of the cross-correlation peak is strongly influenced by the autocorrelation of the neural activity pattern. Interestingly, we found that the correlation between IEG expression levels and maximum calcium activity was higher than that with average calcium activity (**Figures 1E, 1F, S1B and S1C**). Throughout learning, the correlation between IEG expression levels and both average as well as maximum calcium activity were relatively stable with a tendency to increase (**Figure 1F**). Lastly, we quantified the stability of the IEG expression pattern across all 7 training sessions by computing the correlation between the IEG expression pattern in the first recording of each session with the last recording in each session. We found that the IEG expression pattern was relatively stable both within sessions and even across training sessions (**Figures 1G and S1D**).

Given the relatively low correlation of IEG expression levels with neural activity and high stability of IEG expression patterns throughout learning on a population level, we speculated that IEG expression might correlate with a learning-related change in neural activity over the course of the training paradigm. To test this, we first quantified learning-related changes in calcium activity in CA1. We found that a large fraction of the activity was driven by the stimuli used in the 2AFC task. Averaging the responses over all training sessions, we found that about 79% ± 1% (mean ± SEM) in CA1 were significantly activated by at least one of the four stimuli in at least one timepoint (grating: 1.4%; tone: 42.1%; reward: 43.3%; puff: 54.0%; p < 0.01, Student’s t-test over trials) (**Figure 2A**). Many of the tone responsive neurons responded differentially to the 6 kHz and the 11 kHz tones (the average selectivity of the top 10% of tone responsive neurons across timepoints was 0.46, i.e. they responded nearly three times stronger to the preferred tone than the non-preferred (see Methods)). In any given timepoint, on average 54% ± 2% (mean ± SEM) of CA1 neurons were active in the task (we quantified this as the percentage of neurons with at least one calcium transient in a 25 minutes time window to make this value comparable to previous reports (Ziv et al., 2013)). The fraction of active neurons remained constant throughout training (**Figure 2B**) and was lower than the percentage of active neurons in CA1 when mice are navigating a linear virtual tunnel (94% ± 1% (mean ± SEM)). Both the percentage of neurons active in the 2AFC task and the virtual navigation task were higher than the percentage of active neurons in CA1 during exploration of a physical environment previously reported using single-photon calcium imaging (31% ± 1%) (Ziv et al., 2013). Note, however, that at least part of the reason for this lower fraction of active neurons is the lower sensitivity of the single-photon measurements. Thus, the 2AFC paradigm engaged a substantial fraction of CA1 neurons, but as in the case of free exploration the representation was partially dynamic between subsequent timepoints (Ziv et al., 2013). We next quantified how the average population responses to the four stimuli changed from early in learning (timepoint 2; we used timepoint 2 as 2 of the 9 mice did not yet perform the task on timepoint 1) to late in learning (timepoint 7). We found that grating, air puff, and reward responses remained constant, or decreased slightly. The tone responses, however, increased systematically to more than twofold over the course of learning (**Figures 2C and 2D**). Thus, with learning there was a selective increase in the responses to the reward-predicting tone cue.

**Figure 2.**
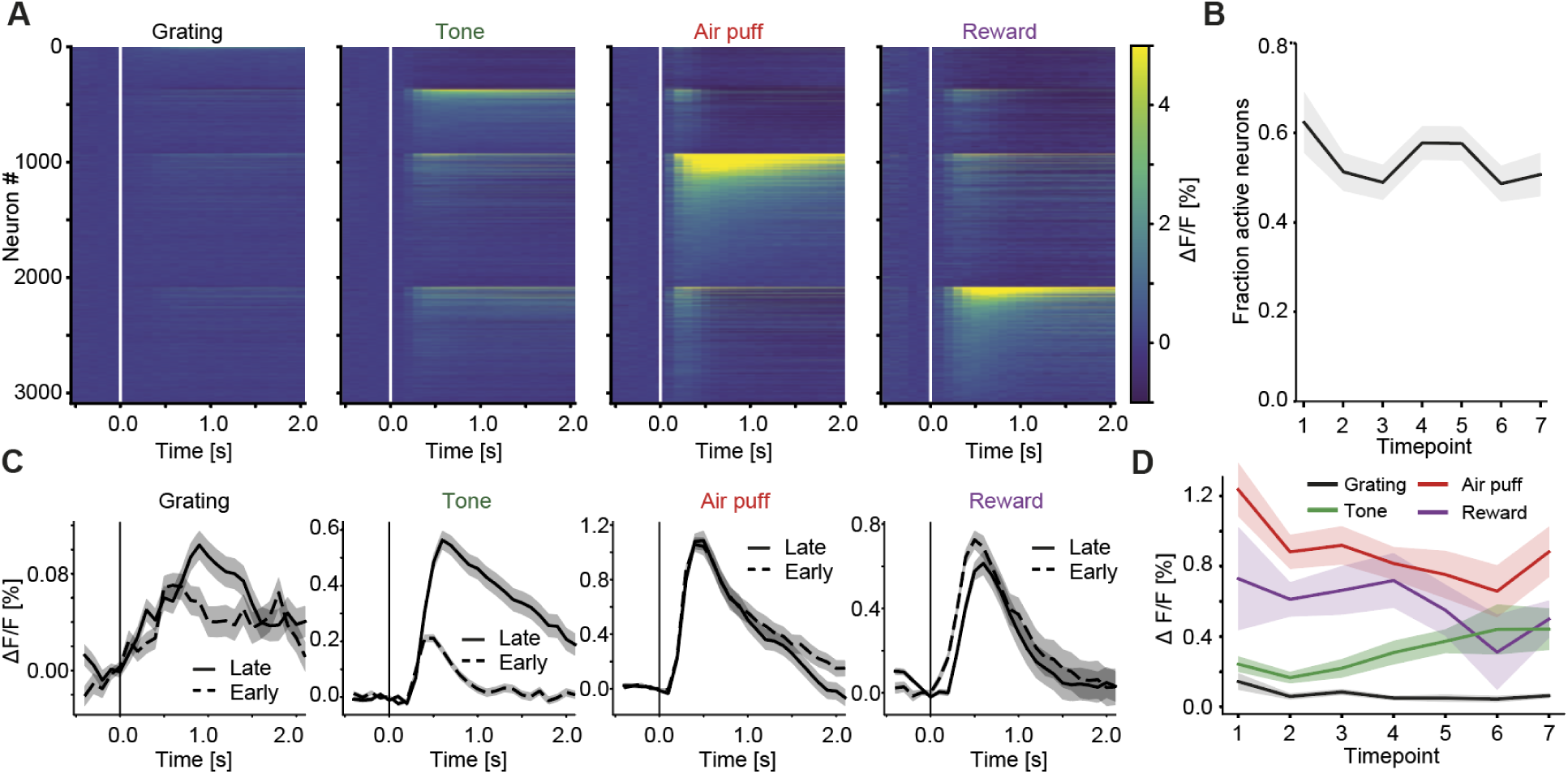
Responses to the task relevant tone stimulus selectively increase with learning. (**A**) Trial-averaged stimulus-evoked activity in all recorded CA1 neurons. Neurons are sorted first by the stimulus they exhibit the strongest response to (sequentially: grating, tone, air puff, and reward) and subsequently strength of response to that stimulus, strongest to weakest. Sorting in all four panels is identical. Data shown are an average over all timepoints. Vertical white lines mark stimulus onset. (**B**) Fraction of active neurons (mean ± SEM) as a function of timepoint. (**C**) Average population responses to the four stimuli in the task early in learning (timepoint 2; dashed lines) and late in learning (timepoint 7; solid lines). The only responses that increase with learning are the tone responses. Shading indicates SEM across neurons (timepoint 2: n = 3090; timepoint 7: n = 2824, data from one animal were discarded on timepoint 7 due to motion artefacts, see Methods). (**D**) Average population responses to the four task stimuli as a function of timepoint. Shading indicates SEM across mice (n = 9). The average tone response is significantly higher during timepoints 5 and 7 than timepoint 2 (9 mice, p < 0.05, Wilcoxon rank-sum test).

Given that responses to the tone selectively increased with learning, it is possible that increased tone responses are predictive of a subsequent correct choice of the mouse. To test whether the strength of the tone response correlated with the choice of the animal and whether this effect is specific to the highest tone responsive neurons, we split the responses of the 10% of neurons with the highest tone responses and those of the remaining population of neurons late in learning (timepoint 7) by trials in which the mouse’s choice was correct or incorrect. The tone responses of high tone responsive neurons were significantly larger in correct trials, while the response of the remaining population was unchanged (**Figure 3A**). To quantify how well we could predict the choice of the mouse based on the activity of the 10% most tone responsive neurons, we trained a binary classification model to predict correct versus incorrect trials based on neural activity as a function of time in trial (see Methods, **Figure 3B**). We found that the activity of the 10% most tone responsive neurons was a better predictor of behavioral choice immediately following the onset of the tone than the remainder of the population. To test whether the correlation between tone response in the 10% most tone responsive neurons and behavioral choice of the mouse developed with learning, we quantified the accuracy of our binary classifier as a function of timepoint in learning (**Figure 3C**). Accuracy was above chance already in session 1 and increased with learning (p = 0.009, R^2^ = 0.12, 7 mice, linear trend analysis). Taken together, these data demonstrate that hippocampal region CA1 is activated by the task in a way that is predictive of the subsequent behavioral choice, and that this activation changes in a learning dependent manner to increase the responses to the reward-predicting tone cue.

**Figure 3.**
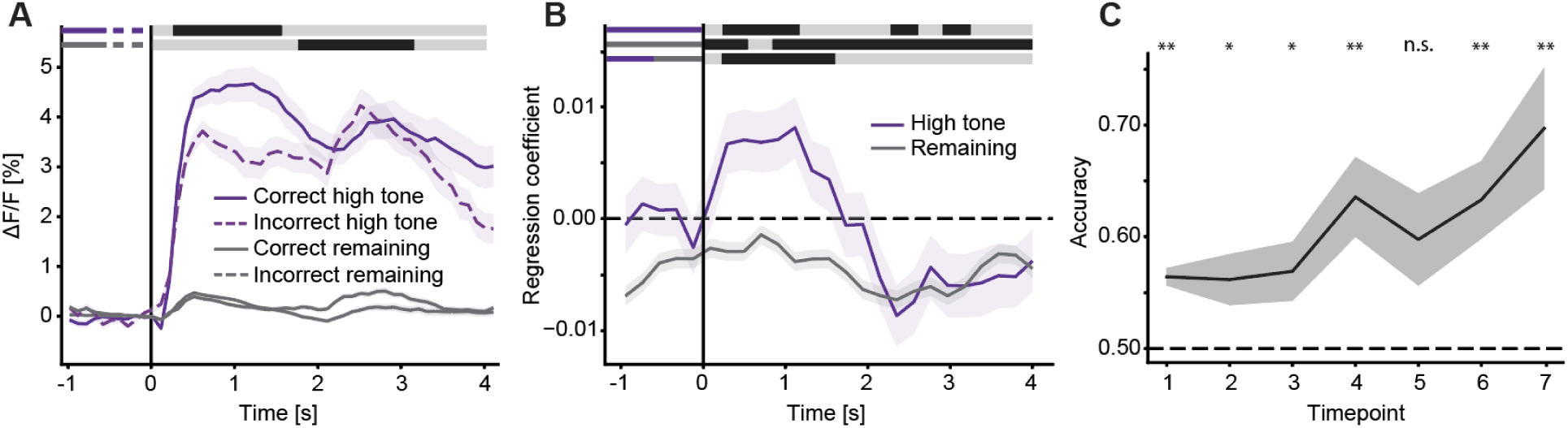
Strength of tone response is predictive of the mouse’s choice. (**A**) The strength of the tone response of the top 10 % tone responsive neurons (purple) correlates with the subsequent behavioral choice in individual trials (solid: correct trials; dashed: incorrect trials). The activity of the remaining population (gray) is similar in correct and incorrect trials. The data shown are from timepoint 7. Shading indicates SEM across neurons. The responses of the high tone responsive neurons and the remaining population in correct and incorrect trials are compared bin-by-bin (100 ms bins) using a Student’s t-test. Bins with a significant difference (p < 0.05) are marked by a black line above the curves; those without are marked as light gray. Each comparison is marked by a pair of line segments to the left, corresponding in color and line style to the curves that are being compared. (**B**) Average regression coefficient of neural activity against trial outcome as a function of time (in 300 ms bins) for the top 10 % tone responsive neurons (purple) and the remaining population (gray). The top 10 % tone responsive neurons (purple) have higher regression coefficients with trial outcome during the tone presentation than the remaining population of neurons. Shading indicates SEM across neurons. The data in the different curves are compared bin-by-bin (200 ms bins) using a Student’s t-test. Bins with a significant difference (p < 0.05) are marked by a black line above the curves; those without are marked as light gray. The upper two lines indicate comparisons against zero, while bottom line indicates the comparison of the two curves. (**C**) Classification accuracy of a binary classification model trained on neural activity in a 1 s window following tone onset (500 ms to 1500 ms) for each timepoint in the task. Horizontal dashed line indicates chance level. Asterisks indicate where accuracy is statistically significant from chance (*: p < 0.05; **: p < 0.01; n.s.: p > 0.05, Student’s t-test). Shading indicates SEM between mice.

One possible explanation of these learning dependent changes in neural activity is plasticity of the local CA1 circuitry. To test for evidence of local plasticity, we quantified how IEG expression levels correlated with learning-related changes in neural activity. We speculated that the neurons that would undergo the strongest learning-related changes in activity would exhibit increased levels of IEG expression during learning. We selected the 10% of neurons with the highest level of IEG expression mid-training, on timepoint 4, and compared tone responses of these neurons the tone responses of the remaining population of neurons. We found that both neurons with high c-Fos and high Arc expression levels exhibited larger tone response increases with learning than the remaining population of neurons (**Figure 4A**). Interestingly, we found that this remained true even when we selected the high IEG neurons in early timepoints, prior to learning-related changes in activity or increases in performance in the task. Starting on timepoint 2, neurons with high levels of Arc expression exhibited larger tone response increases with learning than the remaining population of neurons (**Figure 4B**). For c-Fos expression levels, this was already the case starting with timepoint 1. It is possible that the ability to predict tone response increases based on IEG expression levels is simply a consequence of the fact that IEG expression correlates with features of the neural activity patterns. To test for this, we compared the predictive power of IEG expression levels to that of other features of the neural responses (average and maximum activity, responses to gratings, rewards and air puffs) early in learning by linearly regressing a separation score of the tone increase (d-prime: tone increase normalized by the between-trial variability) to these features. Among the features tested, we found that early Arc and c-Fos expression levels were the best predictors of learning-related tone response increases (**Figure 4C**). Lastly, we tested whether IEG expression levels simply correlate with high tone responses early in learning. We found that high IEG expressing neurons selected on most timepoints had tone responses early in learning comparable to those of the remaining population (**Figure S2**). In sum, these data show that IEG expression levels early in learning correlate with subsequent learning-related changes in tone responses better than would be predicted simply from the correlation between activity patterns and IEG expression. This effect could be explained if increased IEG expression levels would poise neurons for plasticity. Arc protein synthesized prior to learning, for example, has been hypothesized to help consolidate the difference between potentiated and non-potentiated synapses (Minatohara et al., 2015).

**Figure 4.**
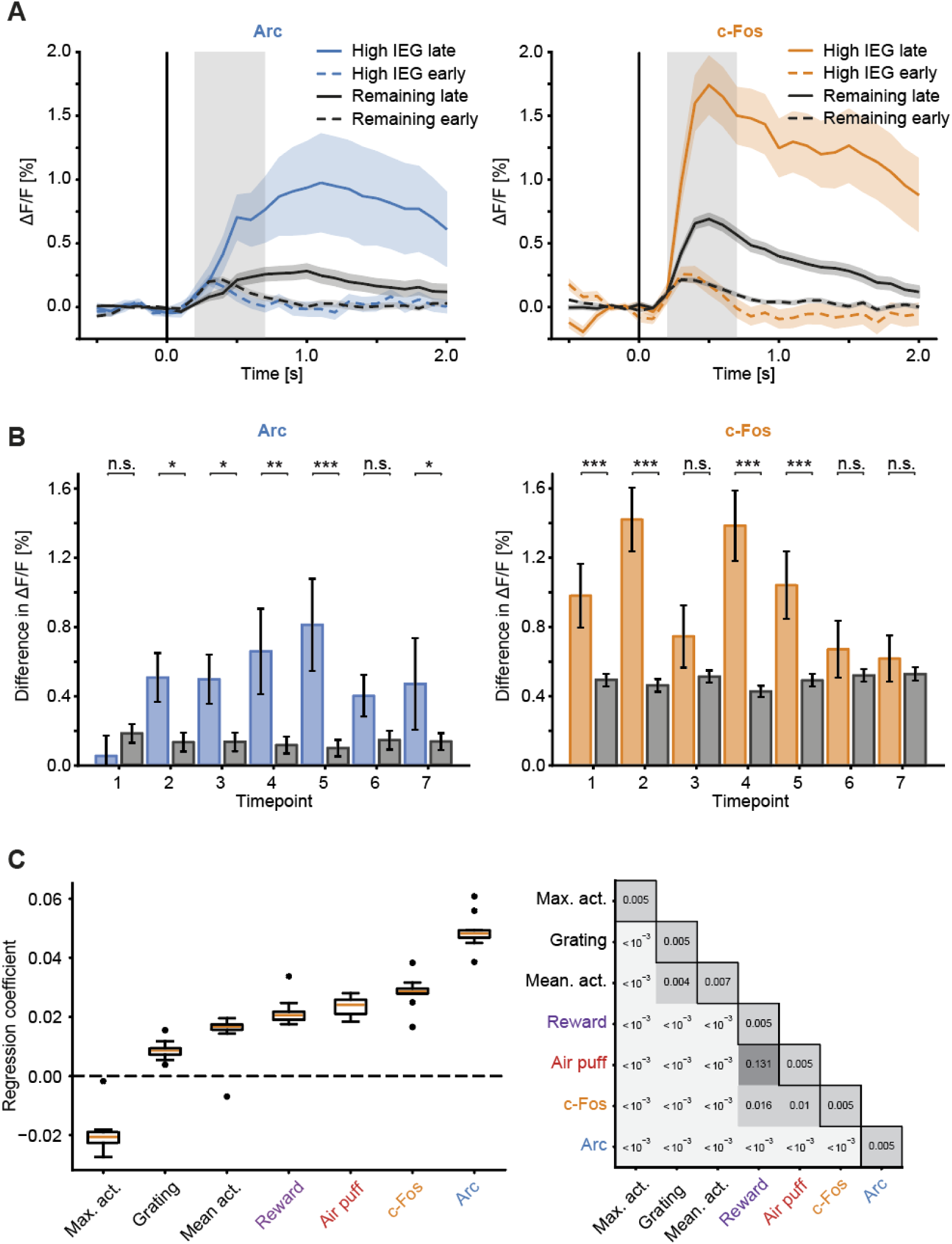
IEG expression levels early in learning correlate with tone response increases during learning. (**A**) Average population responses to the tone onset early (timepoint 2, dashed) and late (timepoint 7, solid) during learning for the top 10% Arc (left) and c-Fos (right) expressing neurons selected on timepoint 4. Note, tone responses of the high Arc and c-Fos-expressing neurons increase more than in the rest of the population. Shading indicates SEM across neurons (high Arc: 128 neurons, remaining: 1143 neurons; high c-Fos: 182 neurons, remaining: 1637 neurons). Gray shading marks the time window used for the quantification shown in **B**. (**B**) Quantification of the effect shown in **A** as a function of timepoint. Blue (orange) bars indicate the difference in tone-evoked activity for high Arc (high c-Fos) expressing neurons between the early (timepoint 2) and late (timepoint 7) timepoints as a function of the timepoint in which the high IEG expressing neurons were selected. Gray bars show the corresponding increase in tone responses in the remaining population of neurons. Error bars indicate SEM across neurons. Asterisks indicate where difference in tone response of high Arc and high c-Fos neurons is different from that of the remaining population of neurons (*: p < 0.05; **: p < 0.01; ***: p < 0.001 n.s.: p > 0.05, Student’s t-test). (**C**) Left: Box plot of regression coefficient distributions between different features early in learning (timepoint 2) and the difference in tone response between timepoints 2 and 7. Orange lines mark the median, boxes the quartiles of the coefficient distributions; whiskers mark the range and black dots outliers. Note, we do not include the early tone responses in this analysis here as these are trivially anticorrelated with changes between early and late tone responses. Right: Significance of all comparisons of the data shown on the left. Off-diagonal elements are the p-values for the comparison of the distributions (Wilcoxon rank-sum test), and the diagonal elements p-values for the difference from 0 for each distribution (Wilcoxon signed-rank test).

## DISCUSSION

The expression of the immediate early genes *c-fos* and *Arc* is often interpreted as a marker of neural activity. Here we quantified this relationship directly in region CA1 of the hippocampus and do indeed find a positive correlation between IEG expression levels and average neural activity (**Figure 1E**). However, the correlation IEG expression levels is higher with maximum activity than with average activity (**Figure 1F**). Given that plasticity is most strongly induced by bursts of activity (Paulsen and Sejnowski, 2000), this would suggest a correlation between IEG expression and plasticity. Consistent with this, we find that neurons that exhibit the highest levels of IEG expression also exhibit the strongest learning-related changes of neural activity (**Figure 4A**). As would be expected from the high level of stability of the IEG expression pattern across learning (**Figure 1G**), this remains true if we select the highest IEG expressing neurons early in learning (**Figure 4B**).

When interpreting our results, it should be kept in mind that both the method we use to measure neural activity as well as the method we use to quantify IEG expression levels have important caveats. Calcium indicators change fluorescence monotonically as a function of neural activity, but the transfer function from spikes to fluorescence change is non-linear (Dana et al., 2016). In addition, single spikes are likely not always detected. Our estimation of changes in Arc and c-Fos protein levels rely on quantifying a fusion protein of a GFP and the IEG product that is overexpressed compared to the endogenous Arc and c-Fos protein levels (Steward et al., 2017). There is, however, a strong overlap between post-mortem antibody stains for GFP and Arc in the Arc-GFP mouse line, as well as between GFP and c-Fos in the GFP-c-Fos mouse line (Barth et al., 2004; Okuno et al., 2012; Yassin et al., 2010). In addition, both onset and offset kinematics of the GFP signal we measure may differ from the endogenous IEG expression levels. Onset kinematics could be affected by a post-transcriptional maturation phase for GFP (Tsien, 1998). Another bias could come from changed degradation time constant of the fusion protein, although this does not seem to be the case at least for c-Fos-GFP (Barth et al., 2004). Both the caveats of the IEG expression level and neural activity measurements could contribute to an underestimation of the correlation between IEG expression level and neural activity, but we see no reason why they would bias the results towards finding increased functional plasticity in neurons with high IEG expression levels.

Optogenetic and pharmacogenetic activation of hippocampal neurons transiently expressing IEGs during experience (Denny et al., 2014; Liu et al., 2012; Ramirez et al., 2013) can drive behavior learned during the conditioning task, and their inactivation impairs memory recall. These studies suggest that neurons expressing *c-fos* or *Arc* during learning are an important part of the ensemble responsible for formation and recall of the respective memory. Our finding that the subset of neurons with high levels of c-Fos and Arc are the ones that selectively undergo experience dependent plasticity would explain why these neurons are critical for memory formation. It is possible that with learning an ensemble of CA1 neurons forms that is characterized by increased expression of IEGs and codes for a specific task relevant context, cue, or time. This supports the idea that IEG expression is not primarily a marker of neural activity, but one of experience dependent plasticity.

## SUPPLEMENTARY FIGURES

**Figure S1 (related to Figure 1).**
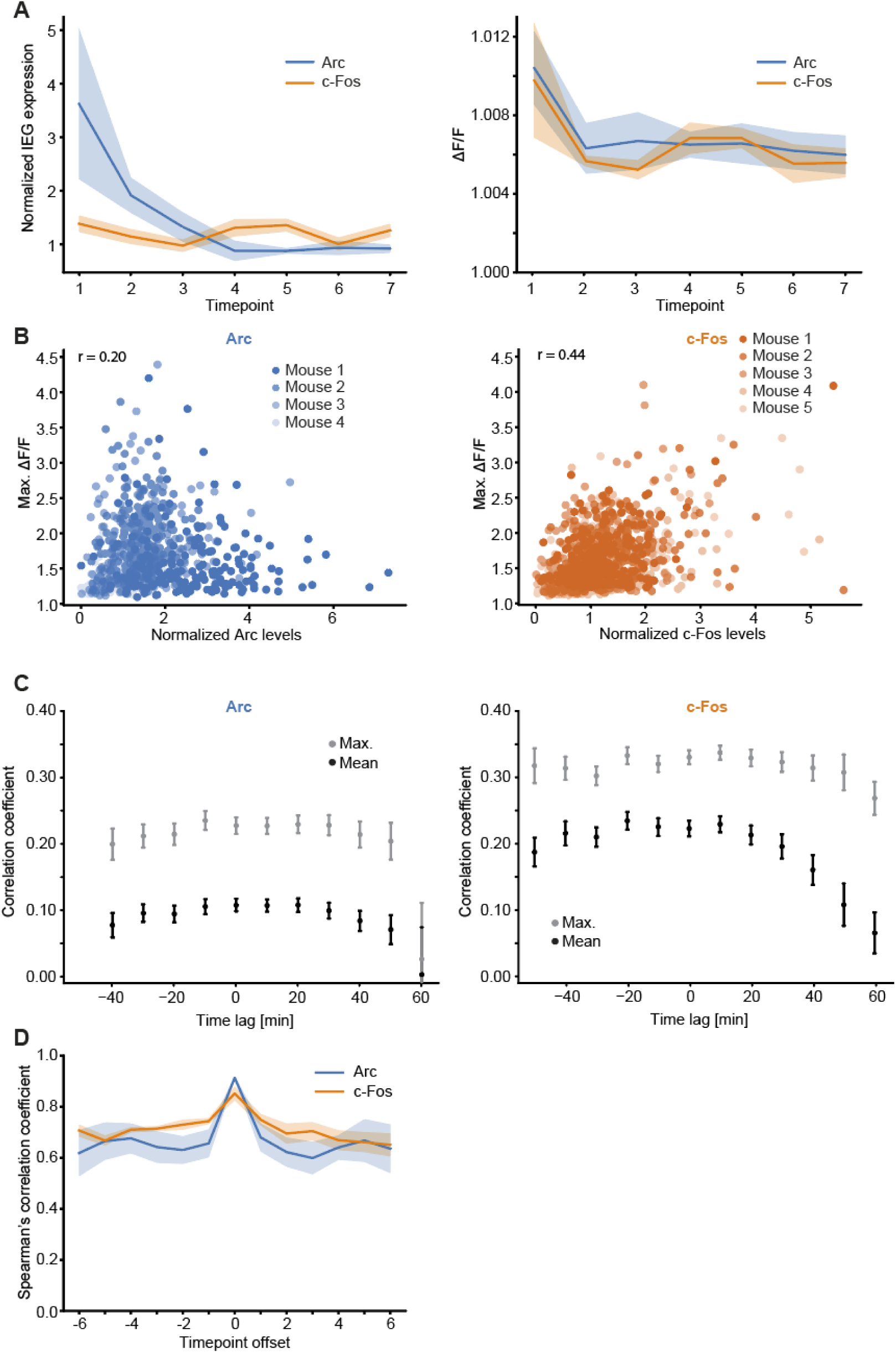
Dynamics of IEG expression and correlation with maximum vs. mean activity. (**A**) Left: Normalized expression of Arc-GFP (blue) and c-Fos GFP (orange) per timepoint. Arc expression levels decrease with time (p = 0.005, R^2^ = 0.26, 4 mice, linear trend analysis), while c-Fos expression levels remain stable (p = 0.746, R^2^ = 0.003, 5 mice, linear trend analysis). Right: Average activity of neurons in the same groups as the left panel. Shading indicates SEM across mice (4 Arc-GFP and 5 c-Fos-GFP mice). (**B**) Scatter plot of maximum calcium activity and IEG expression levels recorded 34 to 42 minutes later for Arc (left) and c-Fos (right) averaged across all timepoints per neuron (Arc-GFP: 1271 neurons; c-Fos-GFP: 1819 neurons). Different colors mark data from different animals (4 Arc-GFP mice, 5 c-Fos-GFP mice). r-value is the average Pearson correlation coefficient across mice. (**C**) Average Pearson’s correlation coefficients between IEG expression and mean (black) and maximum (gray) activity as a function of the time lag between the two, for Arc (left) and c-Fos (right). Error bars indicate SEM across samples. (**D**) Spearman correlation between IEG expression recorded in the first and last IEG measurements in each timepoint (as in **Figure 1G**) as a function of timepoint offset, for Arc (blue), and c-Fos (orange). Shading indicates SEM across mice.

**Figure S2 (related to Figure 4).**
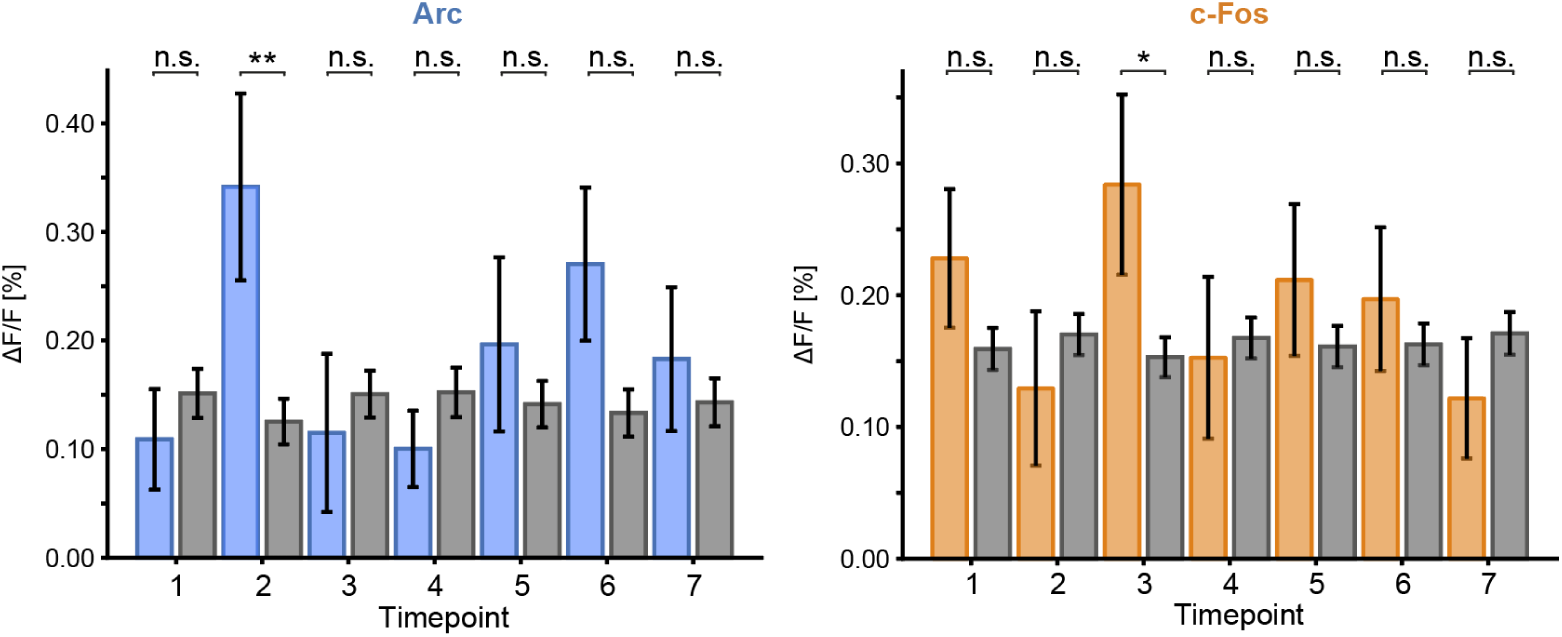
Early tone responses of high IEG expressing neurons. (**A**) Left: The average tone response in timepoint 2 for the top 10% of Arc expressing neurons (blue bars) and the remaining population (gray bars), as a function of timepoint in which the top 10% Arc-expressing neurons are selected. Error bars indicate SEM across neurons. Right: As on the left, but for c-Fos expressing neurons. Asterisks indicate where mean tone response of high Arc and high c-Fos neurons is different from that of the remaining population of neurons (*: p < 0.05; **: p < 0.01; ***: p < 0.001 n.s.: p > 0.05, Student’s t-test).

## METHODS

### Animals and surgery

All animal procedures were approved by and carried out in accordance with guidelines of the Veterinary Department of the Canton Basel-Stadt, Switzerland. We used imaging data from a total of 4 Arc-GFP mice (Okuno et al., 2012) and 5 c-Fos-GFP-mice (Barth et al., 2004), aged 60 to 80 days at the start of the imaging series. All mice were group-housed in a vivarium (light/dark cycle: 12/12 h). No statistical methods were used to predetermine sample sizes. Mice were water-restricted for the duration of the experiment and received water rewards during experiments. Weight of all mice remained above 80% of starting weight. Viral injections and window implantation were performed as previously described (Fiser et al., 2016). Briefly, mice were anesthetized using a mix of fentanyl (0.05 mg/kg), medetomidine (0.5 mg/kg) and midazolam (5 mg/kg) for all surgical procedures. A 3 mm craniotomy was made above either left (in 6 mice) or right (in 3 mice) dorsal hippocampus and posterior parts of cortex were aspirated, and an AAV2/1-Ef1a-NES-jRGECO1a-WPRE (Dana et al., 2016) (titer 1.2 × 10^11^ TU/ml) was injected into hippocampal region CA1. The craniotomy was sealed with a 3 mm cover slip. A titanium head bar was attached to the skull and stabilized with dental cement. Imaging commenced between 23 and 30 days following injection and was done using a custom-built two-photon microscope. Illumination source was an Insight DS laser (Spectra Physics) tuned to a wavelength of either 990 nm or 1030 nm. Imaging was performed using an 8 kHz resonance scanner (Cambridge Technology) resulting in frame rates of 40 Hz at a resolution of 400 × 750 pixels. In addition, we used a piezo-actuator (Physik Instrumente) to move the objective (Nikon 16×, 0.8 NA) in steps of 15 μm between frames to acquire images at four different depths, thus reducing the effective frame rate to 10 Hz.

### Training and experimental design

Mice were handled and accustomed to tubes, similar to the ones used during experiments, in their home-cages five days prior to experiment start by the experimenter. Two days prior to the start of the experiment mice were head-fixed on the setup and randomly rewarded every 20 seconds through one of the two lick spouts to familiarize the mice with the setup. Experimental sessions were 8.3 minutes long, and each experiment consisted of five such sessions. We performed one experiment per day, spaced on average 24 hours apart. On average, mice performed 118 ± 5 trials per day (mean ± standard deviation), during which their performance and activity were recorded. Each trial lasted between 18 and 25 seconds. Trials started with the presentation of one of two oriented gratings selected at random for 2 seconds. Subsequently, one of two tones (6 kHz or 11 kHz) was presented for 4 seconds. Tone identity determined whether the mouse had to lick left or lick right for a water reward. Any lick that occurred during the first 2 seconds of tone presentation was ignored, and the first lick following the 2 second grace period was used to determine the choice of the mouse. Consistent with previous reports (Connor et al., 2010), we found that this grace period during the tone presentation was critical to get the mice to learn the task as they typically sampled both lick spouts in rapid alternation initially and would only focus on licking on one of the two spouts following this initial sampling. A lick on the correct spout would result in the delivery of a water reward, while a lick on the incorrect spout would result in a mild air puff to the neck of the mouse. If the mouse did not lick within the 2 second response window, a mild air puff would be delivered, and the trial ended. The inter trial interval following a correct choice was 14 seconds, and 19 seconds following an incorrect choice or a failure to lick in the response window. To encourage licking, mice were rewarded on 10% of randomly selected trials on the correct lick spout, independent of their behavior. The methods of the experiments in which mice navigated a linear virtual tunnel have been described previously (Fiser et al., 2016).

### Statistics

Non-parametric tests were performed for all analyses (Wilcoxon rank-sum test or Wilcoxon signed-rank test) except where otherwise noted. Linear trend analysis (**Figures 3C and S1A**) was performed using the Scipy linregress function. To quantify the significance of the linear trend we report the R^2^ statistic and the p value of the F statistic.

### Data analysis

No blinding of experimental condition was performed in any of the analyses. Imaging data were full-frame registered using a custom-written software (Leinweber et al., 2014). Neurons were selected manually based on their mean fluorescence or maximum projection of the jRGECO data. The inclusion of the maximum projection biased our selection towards active neurons. Fluorescence traces were calculated as the mean pixel value in each region of interest per frame, and were then median-normalized to calculate *ΔF/F. ΔF/F* traces were filtered as previously described (Mukamel et al., 2009). GFP intensities were calculated as the mean pixel value in each region of interest (ROI). To make IEG measurements comparable across mice, we normalized raw GFP expression values per mouse. We first determined the minimum GFP value across all time points and ROIs per mouse. We then subtracted this minimum value from all GFP expression values and normalized all values by the median of the resulting distribution. This resulted in a distribution of GFP expression values larger than 0 with a median of 1.

For quantification of active neurons (**Figure 2A**), we considered neural activity in the first 25 minutes of each timepoint. We thresholded *ΔF/F* traces by 3.72 times the standard deviation of the noise distribution σ_N_. We estimated σ_N_ as the standard deviation of the lower half of the fluorescence distribution (ΔF/F < median[ΔF/F]) for each cell individually. We then considered a neuron active if there was at least one transient of *ΔF/F* > 3.72 σ_N_ of at least 500 ms duration.

For all plots of stimulus-triggered fluorescence changes (**Figures 2C, 3A and 4A**), fluorescence traces were mean-subtracted in a window 3 to 1 frames (−300 ms to −100 ms) preceding the stimulus onset. Stimulus responses were quantified by the average Δ*F*/*F* in a 5-frame (500 ms, +200 to + 700 ms) window following stimulus onset. Due to uncorrectable brain motion artefacts during imaging, we discarded the imaging data of two mice on the first timepoint (mouse 2 and 9), and of one animal in the last timepoint (mouse 9) for stimulus-evoked activity analysis. In addition, two mice (mouse 1 and 2) did not perform the task in the first timepoint.

Neuronal selectivity to the two tones was defined as

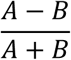

where A and B are the averaged evoked responses in the window described above. We set the selectivity to zero for responses below 0.15% Δ*F*/*F*, to avoid having the mean selectivity be dominated by neurons with tone responses close to zero.

Classification of correct vs. incorrect trials using neural activity (**Figures 3B and 3C**) was performed using a logistic regression model (available in Python’s Scikit-Learn). Separate models were trained for each time bin, on each training session for each mouse. We balanced the labels on each training session by keeping the same number of samples for each class (determined by whichever class had the fewest samples), and we only trained the model if there were at least 8 samples. Each model was trained 20 times on random subsets of half (50%) of the data, and we report the average accuracy on the testing set for each time bin.

Linear regression of the separation score of the tone difference to IEG expression levels and neural activity to different features (**Figure 4C**) was performed using ordinary least squares linear regression (Scikit-Learn). The tone difference was calculated as the difference in average tone-evoked activity in a 500 ms (+200 ms to +700 ms) window in timepoint 7 and timepoint 1. The difference was then normalized by the square root of the sum of the between-trial variances for each tone-evoked response. Data were log-normalized to correct for skewness in the distributions. We then performed 10-fold cross-validation, i.e. trained 10 different regression models on 10 different, independent, training sets and tested them on the remainder of the data. This resulted in 10 point estimates for the regression coefficients, as shown in **Figure 4C**.

### Code availability

All imaging and image processing code can be found online at https://sourceforge.net/projects/iris-scanning/ (IRIS, imaging software package) and https://sourceforge.net/p/iris-scanning/calliope/ (Calliope, image processing software package). The Python code used for all data analysis is available at https://data.fmi.ch.

### Data availability

The data that support the findings of this study are available at https://data.fmi.ch.

## ACKNOWLEDGEMENTS

We thank the members of the Keller lab for helpful discussion and comments on earlier versions of this manuscript. We thank Daniela Gerosa-Erni for production of the AAV vectors, and the members of the Keller lab for technical support. This work was supported by the Swiss National Science Foundation (DM, AF, GBK), the Novartis Research Foundation (DM, AF, GBK), the Human Frontier Science Program (GBK), and the JSPS-KAKENHI grants (HO and HB).

## AUTHOR CONTRIBUTIONS

D.M. and A.V.P performed the experiments. A.F., D.M. and A.V.P analyzed the data. H.O. and H.B. made the mEGFP-Arc mouse. All authors wrote the manuscript.

